# Enhanced *IGFL1* translation in response to IL-1β is controlled by distinct 3’UTR elements

**DOI:** 10.64898/2026.01.21.700974

**Authors:** Giulia Cardamone, Melanie Flohr, Rebecca Raue, Irina Bode, Sofie P. Meyer, Sven Hauns, Rolf Backofen, Tobias Schmid

**Affiliations:** Institute of Biochemistry I, Faculty of Medicine, Goethe University Frankfurt, Frankfurt am Main, Germany; Chair of Bioinformatics, University of Freiburg, Freiburg, Germany; BIOSS Centre for Biological Signaling Studies, Cluster of Excellence, University of Freiburg, Freiburg, Germany; German Cancer Consortium (DKTK), Partner Site Frankfurt, Frankfurt, Germany

## Abstract

Translation is a crucial regulatory mechanism involved in several diseases, including cancer, where pro-inflammatory conditions within the microenvironment have been shown to modulate the translation of specific mRNAs. In the present study, we focused on the regulation of insulin growth factor-like family member 1 (*IGFL1*) in MCF7 breast cancer cells in response to pro-inflammatory IL-1β and observed an induction of both transcription and translation. We characterized the 3’ untranslated region as regulatory hub for the post-transcriptional regulation and identified a distinct G-rich region to confer the IL-1β-dependent translational increase. Our study therefore provides new insights into the translation regulation of *IGFL1* in the context of an inflammatory tumor microenvironment.

## Introduction

Post-transcriptional processes, including translation regulation, are increasingly recognized as key regulatory mechanisms in several disease contexts, including cancer [1,2] and inflammatory diseases [3,4]. Tumor progression is largely controlled by the tumor microenvironment (TME) [5], and the TME is characterized by infiltrating immune cells, including macrophages, that orchestrate the inflammatory, tumor-promoting properties of the TME [6]. The secretion of cytokines and other mediators within the TME has been demonstrated to affect translation, mainly by modulating pathways affecting mTOR kinase activity [7,8]. Pro-inflammatory cytokines, such as interleukin (IL)-6, IL-1α, and IL-1β, appeared to also affect the translation of specific mRNA [9,10]. Concerning IL-1β, it was demonstrated to directly regulate the translation of *THBD* and *EGR2* mRNAs in A459 human lung adenocarcinoma and MCF7 breast tumor cells, respectively [11,12].

The focus of the present work was the characterization of *IGFL1*, which was previously shown to be involved in breast cancer and in inflammatory diseases [13,14]. The *IGFL1* gene encodes the insulin growth factor (IGF)-like family member 1 (IGFL1) protein, belonging to the IGF-like family, consisting of 4 genes (*IGFL1* to *IGFL4*) and two pseudogenes (*IGFL1P1* and *IGFL1P2*) [15]. *IGFL1* expression was first detected in ovary and spinal cords [15] and was shown to be upregulated in skin from psoriasis patients and in primary keratinocyte cultures treated with tumor necrosis factor (TNF)-α [14]. Furthermore, *IGFL1* expression was detected in various breast cancer cells [13,16]. Concerning *IGFL1* expression regulation, it was found to be modulated by the long non-coding RNA IGF-like family member 2 antisense RNA 1 (*IGFL2-AS1*) [16]. In detail, Wang and colleagues [13] showed that the expression of both *IGFL2-AS1* and *IGFL1* was increased through KLF transcription factor 5 (KLF5) upon TNF-α stimulation in basal-like breast cancer cells and that *IGFL2-AS1* contributed to KLF5-dependent *IGFL1* upregulation. With respect to *IGFL1* post-transcriptional regulation, *IGFL2-AS1* was proposed to act as a sponge for a microRNA (miRNA) binding to the *IGFL1* 3’ untranslated region (UTR) [13].

In this work, we identified IL-1β as a novel stimulus to induce *IGFL1* transcription and translation in MCF7 breast cancer cells and characterized its post-transcriptional regulation to be mediated by a specific, G-rich region within its 3’UTR.

## Materials and methods

### Chemicals

All chemicals were obtained from Sigma-Aldrich (Taufkirchen, Germany), if not stated otherwise. Recombinant human IL-1β was purchased from PeproTech (Hamburg, Germany).

### Cell culture

MCF7 cells were purchased from ATCC-LGC Standards GmbH (Wesel, Germany) and were cultured in RPMI 1640 GlutaMAX medium (Gibco, Dreiech, Germany) supplemented with 10% heat-inactivated fetal calf serum (Capricon Scientific, Ebsdorfergrund, Germany), 1% sodium pyruvate, 100 U/mL penicillin, and 100 μg/mL streptomycin. Cells were grown at 37 °C in a humidified atmosphere with 5% CO_2_.

### RNA isolation, reverse transcription, and quantitative polymerase chain reaction (RT-qPCR)

Total RNA was isolated using TRIzol (Thermo Fisher Scientific, Dreieich, Germany) according to the manufacturer’s instructions. RNA was reverse transcribed with the Maxima First Strand cDNA Synthesis Kit (Thermo Fisher Scientific), and qPCR analyses were accomplished by using PowerUp SYBR Green Master Mix on QuantStudio 3 and 5 PCR Real-Time Systems (Thermo Fisher Scientific). Primers were obtained from Biomers (Ulm, Germany) and are listed in S1 Table.

### Polysomal fractionation

MCF7 were subjected to polysomal fractionation as described previously [12]. Briefly, 7.5 x 10^6^ MCF7 cells were seeded in a 15 cm dish, one day prior to polysome fractionation and treated with IL-1β (50 ng/mL) 4 h prior to harvest. Subsequently, cells were incubated with 100 µg/mL cycloheximide (CHX, Carl Roth, Karlsruhe, Germany) for 10 min, washed with PBS/CHX (100 µg/mL), and lysed in 750 µL polysome lysis buffer (140 mM KCl, 20 mM Tris-HCl pH 8.0, 5 mM MgCl_2_, 0.5% NP-40, 0.5 mg/mL heparin, 1 mM DTT, 100 U/mL RNasin (Promega, Walldorf, Germany), 100 µg/mL CHX). After pelleting the cell debris (16,000 x g, 5 min, 4 °C), 600 µL of the cell lysates were layered onto 11 mL of 10–50% continuous sucrose gradients. Gradients were centrifuged at 35,000 rpm for 2 h at 4 °C without brake using a SW40 rotor in an Optima L-90K Ultracentrifuge (Beckman Coulter, Brea, CA, USA). The gradient was collected into 1-mL fractions using a Gradient Station (BioComp Instruments, Fredericton, Canada), and UV absorbance was measured at 254 nm. RNA was precipitated by adding 1/10 volume of sodium acetate (3 M) and 1 volume of isopropyl alcohol. RNA was further purified using the NucleoSpin RNA kit (Macherey-Nagel, Düren, Germany) according to the manufacturer’s instructions. RNA obtained from polysomal fractions was reverse transcribed and quantified by qPCR as described above.

### Plasmid constructs

The psiCHECK-2 vector (Promega) was digested with *Nhe*I or with *Asi*SI and *Not*I (New England Biolabs, Frankfurt am Main, Germany) to insert either the *IGFL1* 5’UTR or *IGFL1* 3’UTR and its truncated versions, respectively. All the inserts were amplified from human cDNA using appropriate PCR primer couples and the PCR product was purified using the NucleoSpin Gel and PCR Clean-up kit (Macherey-Nagel). The fragments were inserted into the linearized vector with the In-Fusion HD Cloning Kit (Takara Bio Europe, Saint-Germain-en-Laye, France) according to the manufacturer’s protocol. The construct 1-413 Δ207-247 was obtained by a cloning reaction of two fragments. All the other constructs containing either a deletion or a point mutation were obtained by site-directed mutagenesis, by means of the QuikChange II Site-Directed Mutagenesis Kit (Agilent Technologies, Waldbronn, Germany) following the manufacturer’s instructions.

All plasmids were purified using the NucleoSpin plasmid kit (Macherey–Nagel) and were verified by Sanger sequencing (SeqLab-Microsynth, Göttingen, Germany). All primers are listed in S1 Table.

### Transient transfection and luciferase reporter assay

5 x 10^4^ MCF7 cells were seeded in 24-well plates 24 h prior to transfection and were transfected with 250 ng of plasmid using jetPRIME (Polyplus, Illkirch-Graffenstade, France), as described by the manufacturer.

Cells were harvested 48 h after transfection. IL-1β-treated cells were treated with 50 ng/mL IL-1β during the last 4 h. Cells were lysed in 100 µl Passive Lysis Buffer (Promega) and the activities of *firefly*/*renilla* luciferase were measured by using the Dual-Luciferase Reporter Assay System (Promega) on a Spark multimode microplate reader (Tecan, Männedorf, Switzerland). *Renilla* luciferase activity was normalized to the corresponding *firefly* luciferase activity, that served as internal transfection control.

### RNA secondary structure predictions

RNA secondary structures were predicted using RNAfold [17] with default parameters and subsequently drawn with the RNA visualization tool VARNA [18].

### Statistical analyses

Statistical analyses were performed with GraphPad Prism v10.6.0 (GraphPad Software, San Diego, CA, USA). Data are reported as means ± SEM of at least three independent experiments. Normal distribution was assessed using the D’Agostino & Pearson test, Anderson-Darling test, Shapiro-Wilk test, and Kolmogorov-Smirnov test. If residuals were assumed to be not normally distributed based on all four tests, data were log-transformed before statistical testing. Statistically significant differences were calculated using paired t-test or two-way ANOVA (either with Tukey’s, Šídák’s, or Dunnett’s multiple comparisons test).

## Results

### IL-1β induces *IGFL1* expression transcriptionally and translationally

*IGFL1* mRNA expression was shown earlier to be induced by TNF-α in basal-like breast cancer cells [13]. To assess, if other pro-inflammatory mediators of the tumor microenvironment might affect *IGFL1* expression in breast tumor cells as well, we stimulated MCF7 breast cancer cells with IL-1β (50 ng/mL, 4 h). Similar to the response to TNF-α, *IGFL1* mRNA expression significantly increased upon IL-1β stimulation (Fig 1A). Since we previously observed that IL-1β commonly affects translational processes in a macrophage-generated tumor microenvironment [12], we next assessed the impact of IL-1β on *IGFL1* translation by polysome profiling. While IL-1β treatment did not alter global translation (Fig 1B) or translation of glyceraldehyde-3-phosphate dehydrogenase (*GAPDH*) mRNA as compared to untreated controls, *IGFL1* mRNA distribution in response to IL-1β treatment significantly increased in the late polysomal fractions (8 and 9), which contain the efficiently translated mRNAs (Fig 1C). Concomitantly, *IGFL1* mRNA abundance decreased in the sub-polysomal and early polysomal fractions.

**Fig 1.**
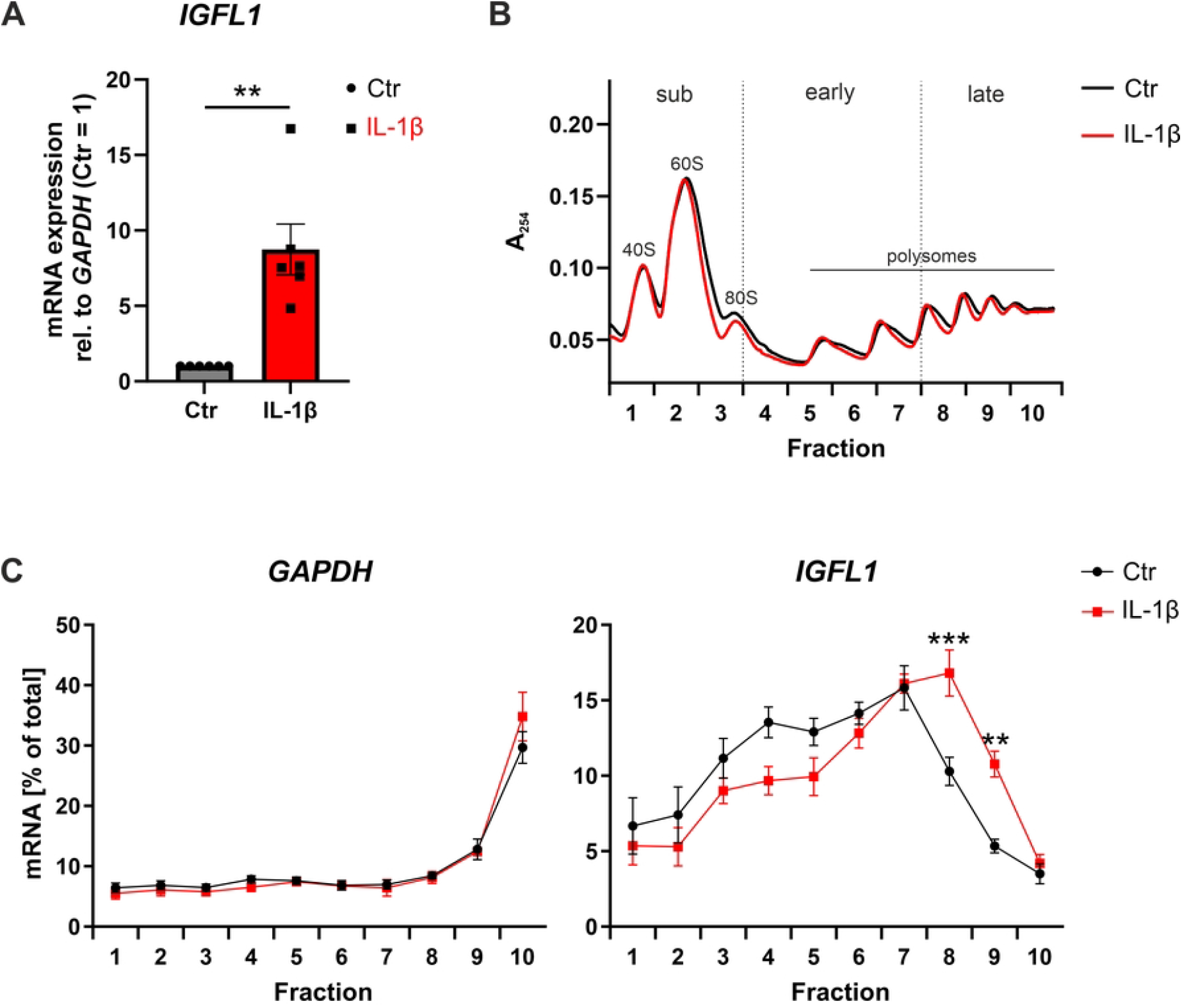
Effect of IL-1β on *IGFL1* mRNA expression and translation. MCF7 cells were treated with IL-1β (50 ng/mL) for 4 h. (**A**) *IGFL1* mRNA expression was measured by RT-qPCR and normalized to *GAPDH* expression (n = 6). (**B, C**) Translational status of *IGFL1* was assessed by polysomal fractionation analysis. (**B**) UV profiles obtained for the sucrose gradients during fractionation for the untreated control and IL-1β-treated cells are shown (representative tracks of four independent experiments). Peaks corresponding to 40S, 60S, and 80S ribosomes and polysomes are depicted. Sub-polysomal (sub), early, and late polysomal fractions are shown. (**C**) *GAPDH* (*left*) and *IGFL1* mRNA (*right*) distribution across the gradients was analyzed by RT-qPCR (n = 4). All data were presented as means ± SEM and statistically analyzed using paired t-test (**A**) or two-way ANOVA with Šídák’s multiple comparisons test; ** *p* < 0.01, *** *p* < 0.001 compared to respective untreated controls.

Thus, IL-1β apparently not only increases *IGFL1* mRNA expression, but also enhances *IGFL1* translation efficiency.

### IL-1β-dependent changes in *IGFL1* translation are controlled by a distinct part of the 3’UTR

Since translation regulatory mechanisms commonly involve the 5’UTR of transcripts, but can also be affected by the 3’UTRs [19,20], we asked if the UTRs might contribute to the IL-1β-dependent translational regulation of *IGFL1* as well. To this end, we cloned the *IGFL1* 5’ and 3’UTRs into the psiCHECK-2 reporter vector up- and downstream of the *renilla* luciferase coding region, respectively, and transfected the resulting vectors into MCF7 cells. Surprisingly, insertion of the 5’UTR neither altered the luciferase activity compared to the empty vector under control conditions nor in response to IL-1β (Fig 2). In contrast, insertion of the *IGFL1* 3’UTR significantly increased the luciferase activity compared to both the empty vector and the vector including the 5’UTR already under basal conditions. Moreover, the 3’UTR-containing vector was markedly more responsive to IL-1β stimulation.

**Fig 2.**
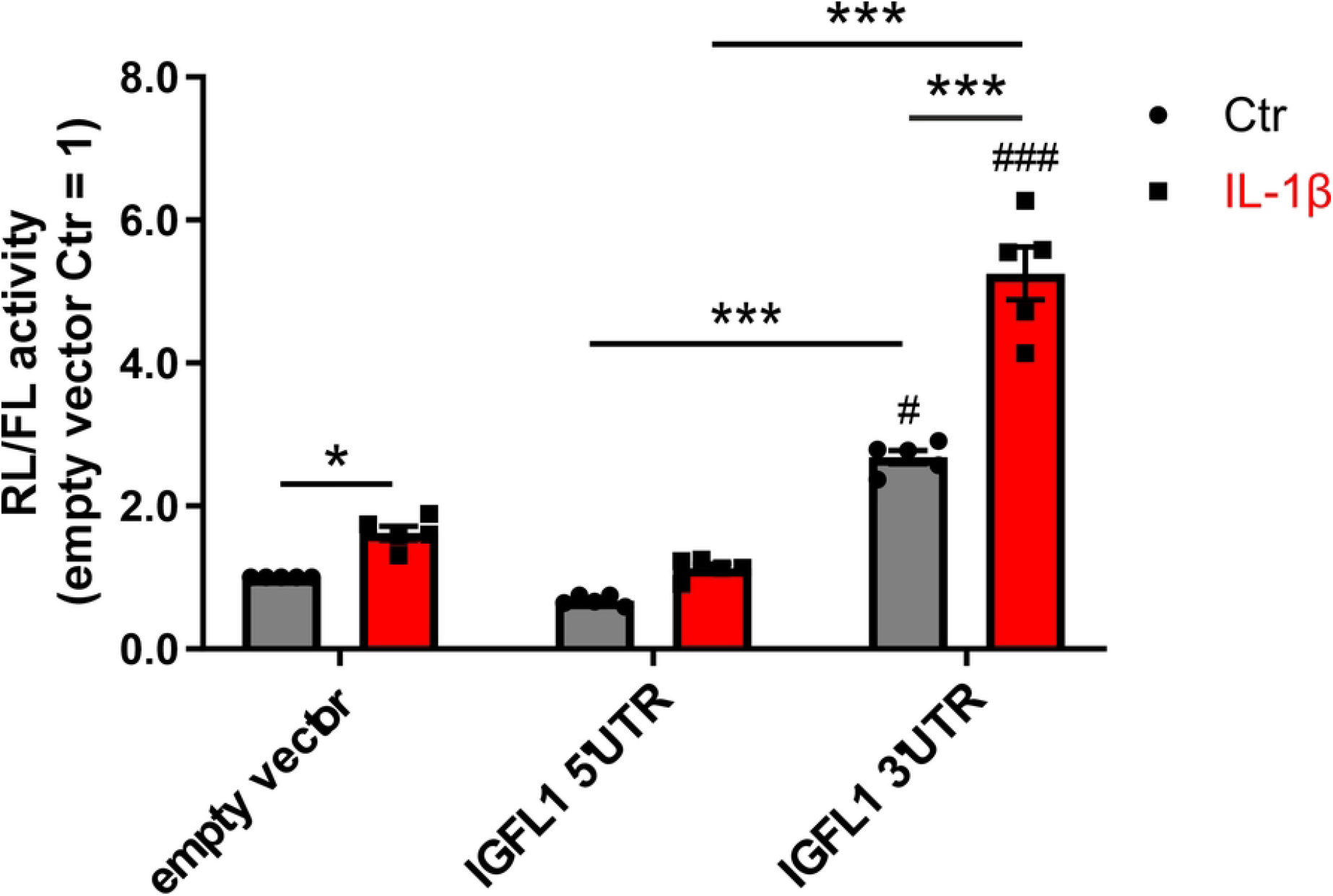
*IGFL1* 3’UTR shows translation-regulatory properties. *IGFL1* 5’UTR and 3’UTR were inserted into the psiCHECK-2 vector upstream and downstream of the *renilla* luciferase coding region, respectively. *Firefly* luciferase served as internal transfection control. MCF7 cells were transfected with the psiCHECK-2 vector (empty vector) or with the psiCHECK-2 vector containing either *IGFL1* 5’UTR or 3’UTR. *Renilla* (RL) and *firefly* luciferase (FL) activities were determined 48 h after transfection with or without treatment with IL-1β (50 ng/mL) during the last 4 h. Data were normalized to the empty vector control, presented as means ± SEM (n = 5), and statistically analyzed using two-way ANOVA with Tukey’s multiple comparisons test; * *p* < 0.05, *** *p* < 0.001;^#^ *p* < 0.05, ^###^ *p* < 0.001 compared to the respective empty vectors.

To get further insights into the role of the 3’UTR in the translational regulation of *IGFL1*, we aimed to identify the exact region within the 3’UTR of *IGFL1* responsible for the IL-1β-induced increased translation. Therefore, we systematically shortened the *IGFL1* 3’UTR within the luciferase vector system. Specifically, we generated constructs representing 75% (1-309), 50% (1-206), and 25% (1-103) of the full length (1-413) *IGFL1* 3’UTR, plus two constructs representing intermediate sizes (1-279 and 1-247) (Fig 3A). Under basal, i.e., unstimulated, conditions (Ctr), the increase in reporter activity appeared to correlate rather directly with the length of the 3’UTR, i.e., the shorter the 3’UTR fragment the lower the respective activity. Conversely, the IL-1β-elicited increase of the 3’UTR-dependent luciferase activity remained comparable to the vector containing the full length 3’UTR for all constructs incorporating at least the first 247 nucleotides of the 3’UTR. Further reduction of the *IGFL1* 3’UTR length to 206 nucleotides led to a substantial drop of the IL-1β response (Fig 3A). These findings suggested that the region between nucleotides 207 and 247 might be relevant for the IL-1β-dependent activation. Modelling of the 2D structure of the entire *IGFL1* 3’UTR reveals that this region encompasses a full, small hairpin structure (nucleotides 211-228) and one arm of a larger hairpin structure (nucleotides 231-284) (Fig 3B). To validate the role of this region and to further rule out a pure length effect, we deleted nucleotides 207-247 in the full length 3’UTR-containing vector (Fig 3B; *green marks*). Indeed, deletion of nucleotides 207-247 (Δ207-247) significantly reduced the IL-1β-induced luciferase activity compared to the full length 3’UTR-bearing vector (Fig 3C). Next, we deleted only nucleotides 235-247, corresponding to a G-rich region embedded within the larger hairpin structure (Fig 3B; *yellow marks*). Strikingly, deletion of this restricted area led to a similar reduction in the IL-1β-induced luciferase activity compared to the deletion of nucleotides 207-247, indicating that this G-rich region might be involved in the IL-1β responsiveness of *IGFL1* translation.

**Fig 3.**
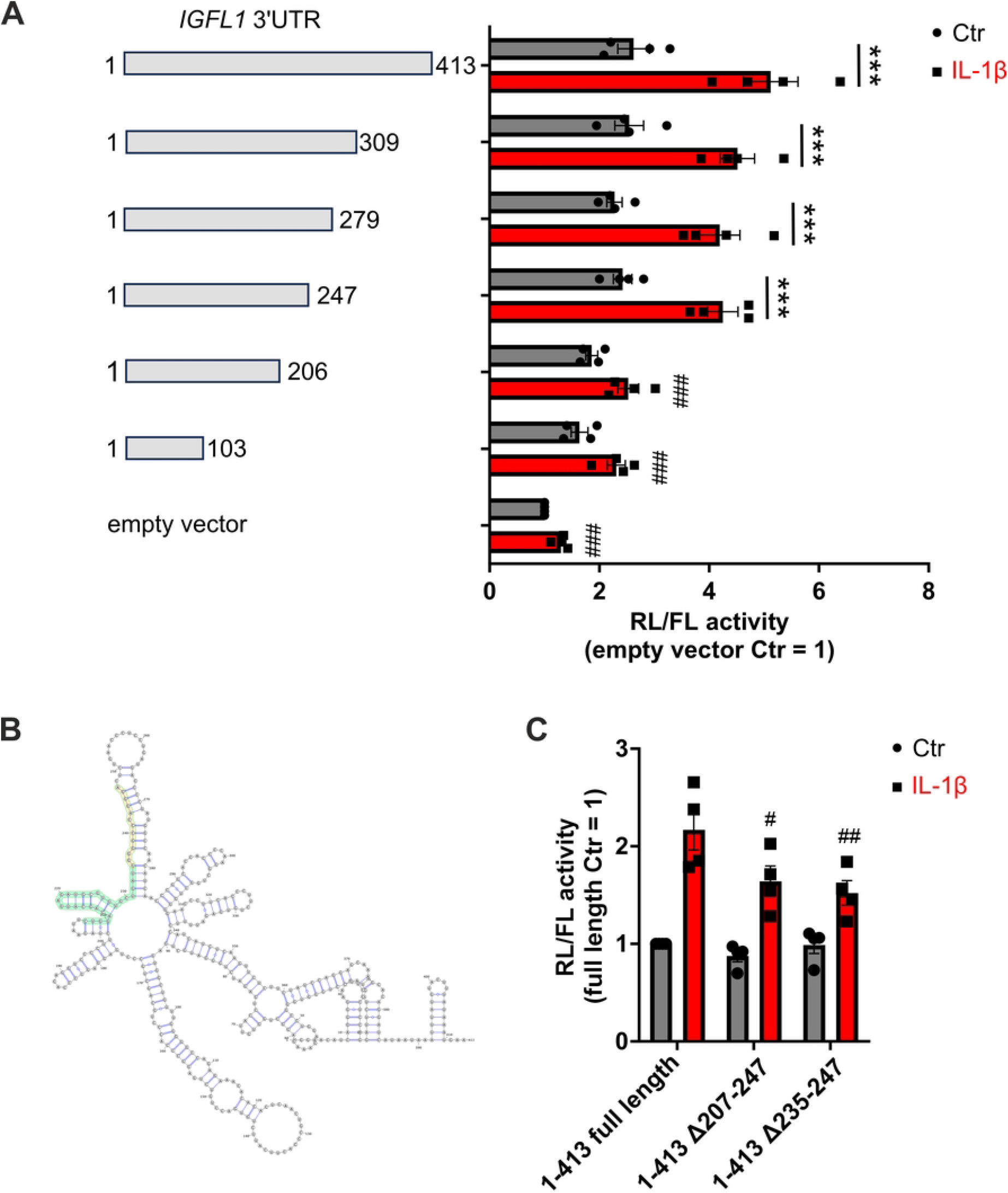
Identification of a translation-regulatory region within the *IGFL1* 3’UTR. MCF7 cells were transfected with the psiCHECK-2 empty vector or with the psiCHECK-2 vector containing either the full length *IGFL1* 3’UTR or parts of it. *Renilla* (RL) and *firefly* luciferase (FL) activities were determined 48 h after the transfection with or without treatment with IL-1β (50 ng/mL) during the last 4 h. (**A**) Schematic representation of the progressive 3’UTR deletion constructs generated in the psiCHECK-2 vector downstream of the *renilla* luciferase coding region and their associated luciferase reporter activities. Data were normalized to the empty vector, presented as means ± SEM (n=4), and statistically analyzed using two-way ANOVA with Tukey’s multiple comparisons test; *** *p* < 0.001; ^###^ *p* < 0.001 compared to the IL-1β-treated full length 3’UTR vector (1-413). (**B**) RNA structure of the full length *IGFL1* 3’UTR (1-413 full length). The depicted structure was generated by VARNA [18]. Nucleotides 207-247 are highlighted in green, nucleotides 235-247 in yellow. (**C**) Luciferase reporter data from MCF7 cells transfected with the psiCHECK-2 vector containing the full length *IGFL1* 3’UTR (1-413 full length), the *IGFL1* 3’UTR deleted of nucleotides 207-247 (1-413 Δ207-247), and the *IGFL1* 3’UTR deleted of nucleotides 235-247 (1-413 Δ235-247). Data were normalized to the full length 3’UTR control vector, presented as means ± SEM (n = 4), and statistically analyzed using two-way ANOVA with Dunnett’s multiple comparisons test; ^#^ *p* < 0.05, ^##^ *p* < 0.01 compared to the IL-1β-treated full length 3’UTR vector.

### Impact of a G-rich region on *IGFL1* translation-regulatory properties

Detailed inspection of the G-rich region revealed that it in fact includes two distinct G-rich motifs, GGGGG at position 235-239 and AGGGA at position 243-247 (Fig 4A). We therefore decided to delete each motif individually (Δ235-239 and Δ243-247) in the full length *IGFL1* 3’UTR construct. Single deletions of the two motifs equally attenuated the IL-1β-responsiveness of the *IGFL1* 3’UTR (Fig 4B), implying an equal importance of these two G-stretches. As both motifs were predicted to be part of the same hairpin structure, we aimed to assess, if the loss of IL-1β responsiveness might be due to a loss of this RNA structural element. Therefore, we introduced point mutations within each motif, predicted to alter the local RNA structure of the *IGFL1* 3’UTR. Specifically, we mutated guanine (G) at position 238 in the lower part of the hairpin structure to a cytosine (C) and G at position 245 to a thymine (T) in the upper part both predicted to lead to massive local structural rearrangement, i.e., a loss of the hairpin between nucleotides 231 and 284. Importantly, both mutations were predicted to leave the structures encompassing nucleotides 1-173 and 284-413 of the full length 3’UTR unaltered, only disrupting the structure in the region in between (Fig 4C; *upper panels*). To unambiguously validate the potential role of this specific structural element, we further introduced corresponding reverse mutations (G238C_C277G; G245T_C269A) predicted to restore the RNA secondary structure to the original *IGFL1* full length 3’UTR structure (Fig 4C; *lower panels*). Introduction of the structure-disrupting mutations indeed attenuated the IL-1β-induced increase in luciferase activity (Fig 4D). Surprisingly though, the reverse mutations did not rescue the IL-1β-induced luciferase activity compared to the structure-disrupting mutations. Since reinstating the *IGFL1* 3’UTR structure did not rescue the IL-1β-responsiveness, the IL-1β-dependent increase in *IGFL1* translation likely does not depend on structural elements within the 3’UTR.

**Fig 4.**
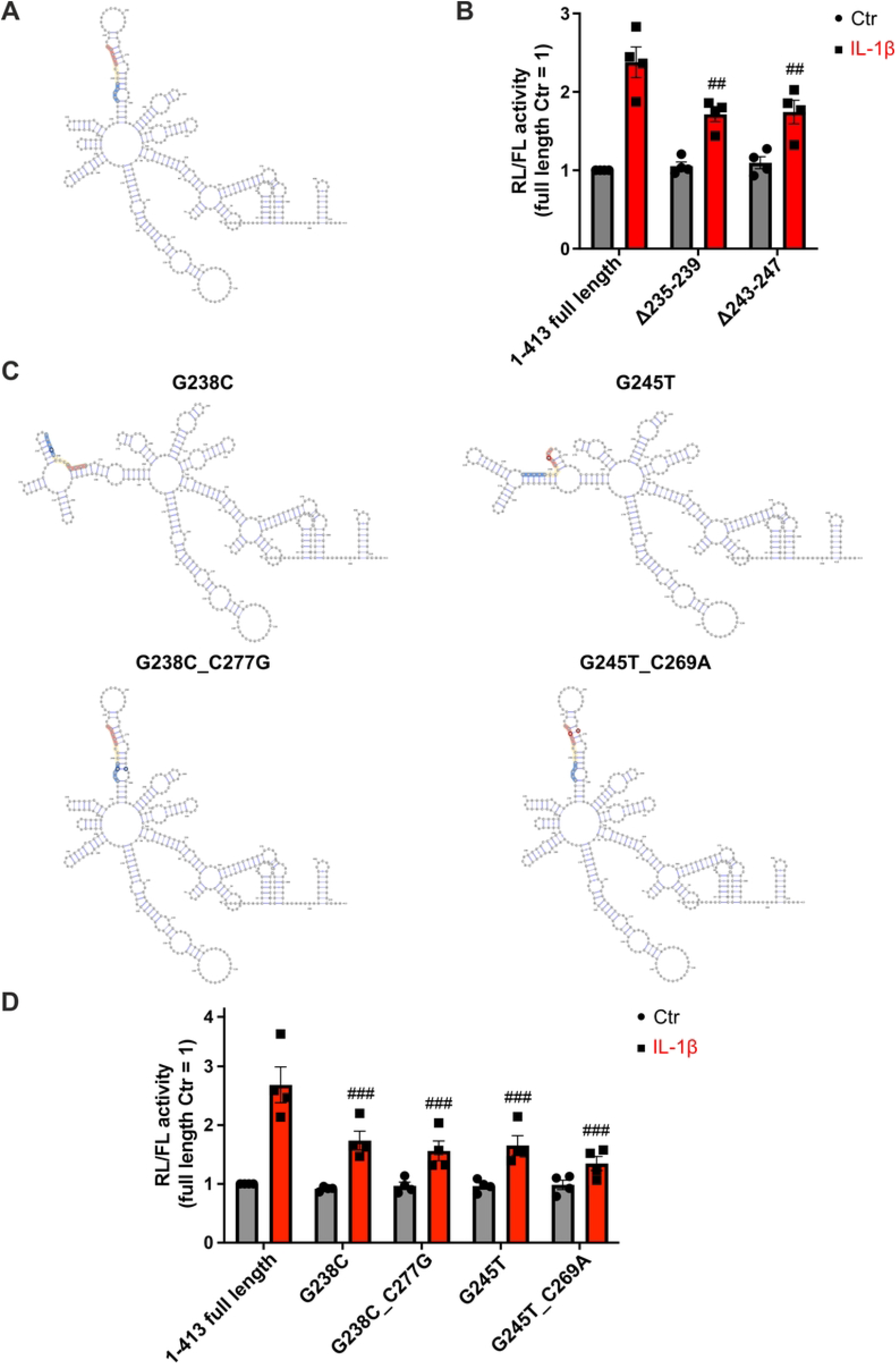
Deletion and mutations in the translation-regulatory region of the *IGFL1* 3’UTR attenuate the IL-1β-dependent increase in *IGFL1* 3’UTR-mediated translation. (**A**) RNA structure of the full length *IGFL1* 3’UTR. Nucleotides 235-247 are highlighted in yellow, nucleotides 235-239 in blue, and nucleotides 243-247 in red. The depicted structure was generated by VARNA [18]. (**B**) MCF7 cells were transfected with the psiCHECK-2 vector containing either the full length *IGFL1* 3’UTR or 3’UTR subjected to deletions. *Renilla* (RL) and *firefly* luciferase (FL) activities were determined 48 h after the transfection with or without treatment with IL-1β (50 ng/mL) during the last 4 h. Data were normalized to the full length 3’UTR control vector, presented as means ± SEM (n = 4), and statistically analyzed using two-way ANOVA with Dunnett’s multiple comparisons test; ^##^ *p* < 0.01 compared to the IL-1β-treated full length 3’UTR vector. (**C**) RNA structures of the full length *IGFL1* 3’UTR with structure-disrupting and respective rescue mutations. The depicted structures were generated by VARNA [18]. Structure-disrupting (G(238)→C) and respective rescue mutations (C(277)→G) are circled in dark blue; structure-disrupting (G(245)→T) and respective rescue mutation (C(269)→A) are circled in red. (**D**) MCF7 cells were transfected with the psiCHECK-2 vector containing either the full length *IGFL1* 3’UTR or 3’UTRs subjected to structure-disrupting and respective rescue mutations. *Renilla* (RL) and *firefly* luciferase (FL) activities were determined 48 h after the transfection with or without treatment with IL-1β (50 ng/mL) during the last 4 h. Data were normalized to the full length 3’UTR control vector, presented as means ± SEM (n = 4), and statistically analyzed using two-way ANOVA with Dunnett’s multiple comparisons test; ^###^ *p* < 0.001 compared to the IL-1β-treated full length 3’UTR vector.

Taken together, our results indicate that the IL-1β-dependent increase in *IGFL1* translation is not mediated by potential RNA structures within the 3’UTR, but rather by regulatory G-rich sequence elements.

## Discussion

In the present study, we demonstrate that IL-1β increases *IGFL1* mRNA transcription and translation in MCF7 breast cancer cells. We further identify the *IGFL1* 3’UTR as regulatory hub for its post-transcriptional regulation and provide evidence that the IL-1β responsiveness of *IGFL1* translation is mediated by a G-rich region within the 3’UTR.

While *IGFL1* has been increasingly recognized as a biomarker in several tumor types as well as in inflammatory diseases [21–25], little is known about its exact function. Similarly, the details of its regulation remain largely elusive. In fact, the pro-inflammatory cytokine TNF-α is the only reported *IGFL1*-inducing stimulus so far, both in the context of psoriasis and in breast cancer cells [13,14]. Here, we extend the panel of *IGFL1* regulating stimuli to IL-1β. Specifically, we observed an IL-1β-dependent induction of *IGFL1* expression in breast cancer cells, both at the level of *IGFL1* mRNA expression and translation. This is in contrast to earlier reports, where *IGFL1* expression was shown to remain not affected by IL-1β in primary keratinocytes [14]. These differences point to a marked tissue- or cell type-specific regulation of *IGFL1*, and, along the same lines, the cell type-specificity of the responses to IL-1β is well-characterized [26–28].

Focusing on the impact of the UTRs in the *IGFL1* induction by IL-1β, we found that the relatively short (33 nucleotides long) 5’UTR did not contribute to the post-transcriptional regulation of *IGFL1*, whereas introduction of the 3’UTR into a reporter vector not only enhanced basal reporter activity, but also showed a marked increase in response to IL-1β. While 5’UTRs are well-established in the regulation of translation initiation, 3’UTRs have also been extensively recognized as important players in the regulation of post-transcriptional processes, including mRNA stability, localization, and translation regulation [29]. 3’UTRs indeed may harbor regulatory sequences including microRNA and RNA-binding protein binding sites as well as structural elements affecting translation regulation [30]. Of note, *IGFL1* was previously shown to be regulated post-transcriptionally by the long non-coding RNA *IGFL2-AS1*, in part by impinging on the 3’UTR of *IGFL1* [13]. Specifically, *IGFL2-AS1* was proposed to compete with *IGFL1* for the binding of miR4795-3p, which is predicted to bind at nucleotide 383 of *IGFL1* 3’UTR. In contrast, we identified the region between nucleotides 207-247 as crucial for the response to IL-1β. Moreover, *IGFL2-AS1* was not or barely detectable in MCF7 cells [13,16]. Instead, we identified two adjacent G-rich motifs (GGGGG and AGGGA) at positions 235-239 and 243-247 of the *IGFL1* 3’UTR, which appeared to contribute to the IL-1β-dependent post-transcriptional induction of *IGFL1*. Indeed, the deletion of the single motifs markedly attenuated the IL-1β-dependent induction of *IGFL1*. Moreover, single nucleotide mutations within each motifs similarly reduced the IL-1β responsiveness. While these single nucleotide mutations were predicted to disrupt the 3’UTR local structure, reversing the local structural changes with respective rescue mutations did not overcome the attenuating effects. This observation rather speaks against the relevance of RNA structures suggesting a sequence-mediated regulatory mechanism in the case of the IL-1β-dependent, post-transcriptional induction of *IGFL1*. It can be envisioned that the two G-rich motifs may be bound by *trans*-acting factors responsible for regulating translation.

In conclusion, we provide evidence that *IGFL1* can be regulated translationally by IL-1β through a 3’UTR-mediated mechanism. In particular, we identify a distinct region within the *IGFL1* 3’UTR, which contains G-rich motifs contributing to the IL-1β-mediated response. It will be interesting to see if the identified mode of regulation is responsible for the cell type-specificity of the *IGFL1* regulation, which might also allow for specific targeting of *IGFL1* expression in tumors or inflammatory diseases.

## Supporting information

**S1 Table. Primers used in this study**.

